# Probing the effects of polysaccharide hydrogel composition on the viability and pro-angiogenic function of human adipose-derived stromal cells

**DOI:** 10.1101/2024.05.10.593603

**Authors:** Fiona E. Serack, Kaylee A. Fennell, Christina Iliopoulos, John T. Walker, John A. Ronald, Brian G. Amsden, David A. Hess, Lauren E. Flynn

**Affiliations:** School of Biomedical Engineering, Faculty of Engineering, The University of Western Ontario, London, Ontario, Canada, N6A 3K7; Department of Chemical & Biochemical Engineering, Faculty of Engineering, The University of Western Ontario, London, Ontario, Canada, N6A 5B9; Department of Anatomy and Cell Biology, Schulich School of Medicine and Dentistry, The University of Western Ontario, London, Ontario, Canada, N6A 5C1; Department of Medical Biophysics, Schulich School of Medicine and Dentistry, The University of Western Ontario, London, Ontario, Canada, N6A 5C1; Robarts Research Institute, The University of Western Ontario, London, Ontario, Canada, N6A 5C1; Department of Chemical Engineering, Faculty of Engineering, Queen’s University, Kingston, Ontario, Canada, K7L 3N6; Department of Physiology and Pharmacology, Schulich School of Medicine and Dentistry, The University of Western Ontario, London, Ontario, Canada, N6A 5C1

**Keywords:** Cell therapy, critical limb ischemia, adipose-derived stromal cells (ASC), hydrogels, chitosan, hyaluronan, vascular regeneration, cell tracking

## Abstract

Cell therapies harnessing the pro-vascular regenerative capacities of mesenchymal stromal cell (MSC) populations, including human adipose-derived stromal cells (hASCs), have generated considerable interest as an emerging treatment strategy for peripheral arterial disease (PAD) and its progression to critical limb ischemia (CLI). There is evidence to support that polysaccharide hydrogels can enhance therapeutic efficacy when applied as minimally-invasive delivery systems to support MSC survival and retention within ischemic tissues. However, there has been limited research to date on the effects of hydrogel composition on the phenotype and function of encapsulated cell populations. Recognizing this knowledge gap, this study compared the pro-angiogenic function of hASCs encapsulated in distinct but similarly-modified natural polysaccharide hydrogels composed of methacrylated glycol chitosan (MGC) and methacrylated hyaluronic acid (MHA). Initial *in vitro* studies confirmed high viability (>85%) of the hASCs following encapsulation and culture in the MGC and MHA hydrogels over 14 days, with a decrease in the cell density observed over time. Moreover, higher levels of a variety of secreted pro-angiogenic and immunomodulatory factors were detected in conditioned media samples collected from the hASCs encapsulated in the MGC-based hydrogels compared to the MHA hydrogels. Subsequent testing focused on comparing hASC delivery within the MGC and MHA hydrogels to saline controls in a femoral artery ligation-induced CLI (FAL-CLI) model in athymic *nu/nu* mice over 28 days. For the *in vivo* studies, the hASCs were engineered to express tdTomato and firefly luciferase to quantitatively compare the efficacy of the two platforms in supporting the localized retention of viable ASCs through longitudinal cell tracking with bioluminescence imaging (BLI). Interestingly, hASC retention was significantly enhanced when the cells were delivered in the MHA hydrogels as compared to the MGC hydrogels or saline. However, laser Doppler perfusion imaging (LDPI) indicated that the restoration of hindlimb perfusion was similar between the treatment groups and controls. These findings were corroborated by endpoint immunofluorescence (IF) staining showing similar levels of CD31^+^ cells in the ligated limbs at 28 days in all groups. Overall, this study demonstrates that enhanced MSC retention may be insufficient to augment vascular regeneration, emphasizing the complexity of designing biomaterials platforms for MSC delivery for therapeutic angiogenesis. In addition, the data points to a potential challenge in approaches that seek to harness the paracrine functionality of MSCs, as strategies that increase the secretion of immunomodulatory factors that can aid in regeneration may also lead to more rapid MSC clearance *in vivo*.

## 1 Introduction

There are currently more than 230 million people worldwide suffering from peripheral arterial disease (PAD), and its prevalence continues to rise due to the increasing frequency of risk factors such as diabetes, obesity, and smoking, as well as the aging population [1]. Critical limb ischemia (CLI) represents the most severe form of PAD and is associated with lower limb amputation rates exceeding 15-20% at 1 year post-diagnosis, and mortality rates greater than 50% by 5 years post-diagnosis [2], [3]. An emerging treatment strategy for PAD/CLI is the delivery of mesenchymal stromal cells (MSCs) to the ischemic tissues, with the goal of stimulating vascular regeneration to counteract the diffuse microcirculatory dysfunction associated with the disease [4]. In particular, adipose-derived stromal cells (ASCs) are a promising cell source due to their relative abundance and accessibility [5], as well as their demonstrated pro-angiogenic and immunomodulatory paracrine functionality [6], [7].

Despite demonstrating promise in pre-clinical models [8], [9], the success of MSC-based therapies has been limited when translated to larger-scale clinical trials in humans [10]. A postulated barrier is the low retention and survival of the cells following injection into the dynamic and highly inflammatory environment within ischemic muscle [9], [11], [12]. Recognizing the need for improved cell delivery strategies, there has been growing interest in the application of biomaterials-based delivery platforms to support localized MSC viability and harness their pro-regenerative functions to improve outcomes in patients with PAD/CLI [12]–[15].

Polysaccharide hydrogels are promising for MSC delivery as they can be designed to be injectable for minimally-invasive delivery, and their high water content and permeability are favorable for cell survival following encapsulation [16]. In particular, chitosan-based biomaterials have been widely explored for wound healing applications, and have been shown to promote angiogenesis [17]–[20] and enhance localized cell retention *in vivo* [13], [21]–[23]. Similarly, hyaluronic acid is a key mediator of wound healing and plays important roles in angiogenesis, making it an attractive base material for the development of pro-angiogenic therapies [24]–[26]. Biomaterial properties such as stiffness, porosity, geometry, and topography are known to influence MSC phenotype and function [16], [27]. However, there has been very limited research on the effects of hydrogel composition on the pro-regenerative function of encapsulated MSCs. Moreover, it is challenging to compare the results of different studies in the literature, due to varied approaches in terms of materials synthesis and functionalization, which can lead to a wide range of physical, chemical, and bioactive properties that could impact the cellular response [28].

With a specific interest in the effects of hydrogel composition on the pro-angiogenic function of encapsulated human ASCs (hASCs), this study compared distinct but similarly-modified natural polysaccharide hydrogels, composed of methacrylated glycol chitosan (MGC) and methacrylated hyaluronic acid (MHA), to advance in the rational design of a cell delivery platform targeting PAD. To more specifically assess hydrogel composition, the MGC and MHA hydrogels were designed to have similar initial Young’s moduli. As an additional comparative group, we included composite hydrogels previously developed by our team for intramuscular cell delivery composed of MGC and a terminally-acrylated triblock copolymer of poly(ethylene glycol) and poly(trimethylene carbonate) (PEG(PTMC-A)_2_) [29] within the *in vitro* studies. Following *in vitro* characterization, the effects of delivery in the MGC or MHA hydrogels on viable hASC retention and *in vivo* vascular regeneration were assessed in a femoral artery ligation-induced critical limb ischemia (FAL-CLI) model in athymic *nu/nu* mice over 28 days. The MGC+PEG(PTMC-A)_2_ group was excluded from the *in vivo* studies for ethical reasons because of previous findings of an adverse inflammatory response when the composite gels were used to deliver hASCs in this model [30]. It was hypothesized that the hydrogel polysaccharide composition would influence the viability and paracrine function of the encapsulated hASCs, ultimately altering their capacity to restore hindlimb perfusion following *in vivo* delivery.

## 2 Methods

### 2.1 Materials

All chemicals and reagents were purchased from Millipore Sigma and used as received, unless otherwise noted.

### 2.2 Polymer synthesis and characterization

MHA synthesis followed a protocol adapted from published methods [31], [32]. In brief, sodium hyaluronate (minimum number average moleculat weight (M_N_)= 100 kDa, Lifecore® Biomedical) was dissolved in deionized water (dH_2_O) at a concentration of 1% (w/v), and the pH was adjusted to 8. Methacrylic anhydride at a molar ratio of 20:1 with respect to the repeating unit of hyaluronic acid was added dropwise over 1 h while the pH was maintained at 8. The reaction was allowed to proceed for 24 h at 4°C with constant stirring. The mixture was then neutralized, and MHA was collected by a series of three precipitations in absolute ethanol at −20°C. The polymer was then dissolved in dH_2_O and the pH was adjusted to 7, followed by dialysis against 0.1 M NaCl using a 6-8 kDa molecular weight cutoff (MWCO) membrane. Purified MHA was characterized using ^1^H NMR spectroscopy in D_2_O at room temperature, showing a degree of methacrylation of 25% (Supplementary Figure S1A).

The synthesis of MGC and PEG(PTMC-A)_2_ were performed according to previously-published protocols [13], [29], [30]. Glycol chitosan (GC) (M_N_ = ∼82 kDa, 85% degree of deacetylation, Wako Chemicals Inc.) was methacrylated via reaction with glycidyl methacrylate. Subsequently, ^1^H NMR spectroscopy in deuterium oxide (D_2_O) at 80°C was used to confirm the degree of methacrylation as 5%, consistent with our previous work [13] (Supplementary Figure S1B). PEG diol (PEG_20,_ MN = 20 kDa) was used to initiate the ring-opening polymerization of trimethylene carbonate (TMC) (LEAPChem). The molecular weight of the polymer was assessed using ^1^H NMR in deuterated DMSO (DMSO-d_6_) on an Inova 600 NMR spectrometer (Varian), confirming the molar ratios of PEG_20_(PTMC_2_)_2_. PEG(PTMC)_2_ was then acrylated via reaction with acryloyl chloride to produce PEG(PTMC-A)_2_ with a high degree of acrylation (85%), consistent with our previous work [13], as confirmed by ^1^H NMR spectroscopy (Supplementary Figure S1C).

### 2.3 Hydrogel preparation and physical characterization

Hydrogels were prepared from the three different polymer systems (MHA, MGC, MGC+PEG(PTMC-A)_2_). Based on previous work [13], [23], a formulation of 1% (w/v) MGC + 4% (w/v) PEG(PTMC-A)_2_ was used for the composite MGC+PEG(PTMC-A)_2_ hydrogels. To generate hydrogels with similar Young’s moduli to control for potential biomechanical effects on cellular and host responses, 3% (w/v) MGC hydrogels and 7.5% (w/v) MHA hydrogels were prepared based on preliminary mechanical testing. All hydrogels were prepared by first dissolving the polymers in phosphate buffered saline (PBS) (Wisent Inc.), using a volume of 95% of the final hydrogel volume. Concentrated solutions of ammonium persulphate (APS) (Bio-Shop) and tetramethylethylenediamine (TEMED) (Bio-Shop) were prepared and added sequentially to the combined mixture with thorough mixing with the pipette tip between each addition, to a final concentration of 5 mM each of APS and TEMED, each making up 2.5% of the final volume.

Hydrogels for the physical characterization studies were prepared in syringe molds using 0.3 mL insulin syringes, and allowed to gel at 37°C for 15 min. Following gelation, 30 μL hydrogels were cut with a razor blade and used to assess the swelling ratio and compressive Young’s moduli. The 30 μL hydrogels were chosen to have a height:diameter ratio greater than 1.5 to minimize the Poisson effect [33]. The height and diameter of the hydrogels were measured using digital calipers, following which the hydrogels were transferred into 12-well plates containing 2 mL PBS/well, and allowed to swell at 37°C for 24 h. The height and diameter of the hydrogels were re-measured following swelling, and the swelling ratio was calculated as the difference in the initial and final volume, divided by the initial volume. A total of 6 MHA and MGC hydrogels and 5 MGC+PEG(PTMC-A)_2_ hydrogels were assessed (n=5-6).

The compressive Young’s moduli of all three hydrogel formulations were assessed using a CellScale MicroTester system (CellScale), following swelling. Hydrated hydrogels were placed in a PBS bath at 37°C and the cantilever was lowered until the upper compression plate made contact with the hydrogel. Hydrogels (n=4 hydrogels/formulation) were compressed to 20% of their initial height for 6 cycles at 0.5 Hz. Stress-strain curves were generated, and the slope of the linear portion was calculated for cycles 3-6 and averaged as the compressive Young’s modulus for each hydrogel.

### 2.4 hASC isolation and culture

Resected human adipose tissue was obtained with informed consent from routine breast or abdominal reduction surgeries conducted at the London Health Sciences Centre in London, ON, Canada, with approval from the Human Research Ethics Board at Western University (HSREB# 105426). Fresh adipose tissue samples were transported to the lab on ice in sterile PBS supplemented with 2% bovine serum albumin (BSA) (Bio-shop) and hASCs were isolated following published methods [34]. The cells were cultured in ASC proliferation medium consisting of Dulbecco’s Modified Eagle Medium:Nutrient Mixture F-12 (DMEM/F12) (Wisent Inc.) supplemented with 10% FBS and 1% penicillin-streptomycin (Gibco™). Media was replaced every 2-3 days and passaging was performed at 80% confluence, with hASCs at passage 3-5 used for all studies. Cell donor information is summarized in Supplementary Table S1.

To enable longitudinal tracking of the delivered cells within the mouse model, hASCs used for the *in vivo* studies were transduced following published protocols to co-express tdTomato (tdT) and firefly luciferase (FLuc2) under the human elongation factor 1-α promoter for constitutive expression (pEF1-alpha-tdT-FLuc2 reporter), with an average transduction efficiency >85% [35]. Following transduction, the cells were cultured as described above and used at passage 4 for the *in vivo* studies.

### 2.5 hASC encapsulation for in vitro studies

To prepare the hydrogels for the *in vitro* studies, dry MHA, MGC, or MGC + PEG(PTMC-A)_2_, were decontaminated under low-intensity UV light in a biosafety cabinet for 30 min. The weight of each polymer was calculated based on a final formulation of 7.5% (w/v) MHA, 3% (w/v) MGC, or 1% (w/v) MGC and 4% (w/v) PEG-(PTMC-A)_2_. The polymers were then dissolved in sterile PBS (75% of the final volume) overnight at room temperature with constant agitation on an orbital shaker at 100 RPM.

hASCs were extracted using trypsin (Wisent Inc.), resuspended at a concentration of 5×10^7^ cells/mL (5X final concentration) in ASC proliferation medium (20% of final volume), and thoroughly combined with the pre-polymer solutions. Concentrated solutions of APS and TEMED were sterile filtered and added sequentially to the combined mixture with thorough mixing with the pipette tip between each addition, to a final concentration of 5 mM each of APS and TEMED. The volumetric ratio of the final solutions was 75% pre-polymer solution : 20% hASC suspension : 2.5% APS : 2.5% TEMED, with a final cell density of 10^6^ cells/mL. Hydrogels for the *in vitro* studies were prepared in syringe molds using 0.3 mL insulin syringes, and allowed to gel at 37°C for 15 min. Following gelation, 10 μL hydrogels were transferred to individual wells of a 12-well plate, and cultured in 2 mL of ASC proliferation medium, with medium changes every 2 days.

### 2.6 Assessment of cell viability in vitro

Live/Dead™ staining was performed at 24 h, 7 d, and 14 d post-encapsulation to assess the viability and density of the hASCs encapsulated within the MHA, MGC, and MGC+PEG(PTMC-A)_2_ hydrogels. Samples were rinsed with PBS, followed by 30 min incubation at 37 °C in a solution composed of 2 μM calcein-AM (Invitrogen™) and 4 μM ethidum homodimer-1 (Invitrogen™) in PBS. Hydrogels were subsequently rinsed with PBS, and imaged using a Zeiss LSM confocal microscope (Zeiss Canada) with a 10X objective. Stitched images were prepared for the entire cross-section of each hydrogel, and quantification of viability and live cell density were performed using ImageJ in 3-5 hydrogels per formulation per timepoint, and repeated with a total of 3 different hASC donors (n=3-5, N=3).

### 2.7 Pro-angiogenic and immunomodulatory paracrine factor release in vitro

The expression levels of a variety of angiogenic and immunomodulatory factors released by the hASCs encapsulated in the different hydrogel formulations were investigated by characterizing conditioned media (CdM) samples using customized Luminex® multiplex assays or an enzyme-linked immunosorbent assay (ELISA). hASCs were encapsulated in the hydrogels and cultured *in vitro* in ASC proliferation medium. At 24 h and 6 d post-encapsulation, samples were rinsed with PBS and transferred into serum-free DMEM/F12 with 1% penicillin-streptomycin. CdM was generated over 24 h and collected at the timepoints of 48 h and 7 d. CdM was collected in sterile 2 mL microcentrifuge tubes and centrifuged at 14,800 xg for 10 min to remove cell debris. The supernatant was transferred to new sterile microcentrifuge tubes and stored at −80°C until needed. The *in vitro* cytokine release studies included 3 replicate hydrogels per formulation / timepoint and were performed a total of 6 times with different hASC donors (n=3, N=6).

The hydrogels were also collected at the 48 h and 7 d timepoints to quantify the total double stranded DNA (dsDNA) content for normalization of the secretome data. Collected hydrogels were rinsed in PBS, snap frozen, and crushed using a plastic pestle. The samples were then digested using proteinase K in buffer ATL (20 mg/mL, >318 mU/mL, Qiagen) overnight at 56°C. DNA was extracted from the samples using a DNeasy® Blood & Tissue Kit (Qiagen), following the manufacturer’s instructions. The total dsDNA content was then quantified using the Quant-iT™ PicoGreen™ assay (Invitrogen™) according to the manufacturer’s instructions, with analysis using a CLARIOstar® spectrophotometer (BMG LABTECH). Controls of unseeded hydrogels were included in the analysis for all three hydrogel formulations. All samples and standards were run in technical triplicate for the dsDNA assay.

A Luminex® multiplex assay (R&D Systems) was used to quantify the concentrations of angiogenin, hepatocyte growth factor (HGF), interleukin-8 (IL-8), monocyte chemoattractant protein-1 (MCP-1), and vascular endothelial growth factor (VEGF) in the CdM from the hASCs cultured in the different hydrogel formulations, according to the manufacturer’s instructions. The samples were analyzed using a MAGPIX™ Multiplex Reader (Bio-Rad), using the xPONENT program with a five-parameter logistic curve-fit. Similarly, an ELISA (DuoSet ELISA, R&D Systems) was used to quantify the concentration of interleukin-6 (IL-6) in the CdM, according to the manufacturer’s instructions. The plates were read using a CLARIOstar® microplate reader at 450 and 540 nm, and the results were analyzed using GraphPad Prism software with a four-point logistic curve-fit. Protein concentrations were determined based on comparison to the standard curves and normalized to the dsDNA content measured in each sample. Controls of unseeded hydrogels were included in the analyses for all three hydrogel formulations.

### 2.8 Pro-angiogenic and immunomodulatory gene expression in vitro

A follow-up study was performed to assess the gene expression levels of the various angiogenic and immunomodulatory factors included in the secretome analyses in the encapsulated hASCs after 7 days in culture, with hASCs cultured on tissue culture polystyrene (TCPS) included as a control. Hydrogel samples were collected, rinsed with PBS, and snap-frozen in PureZOL. Total cellular RNA was isolated using PureZOL™ RNA Isolation Reagent (Bio-Rad) and the RNeasy® Micro Kit (Qiagen), according to the manufacturer’s instructions. Isolated RNA was subsequently dissolved in 3M sodium acetate and reprecipitated using absolute ethanol. A NanoDrop™ (ThermoFisher) was used to quantify the concentration and purity of the isolated RNA. cDNA was generated from the isolated RNA using an iScript™ cDNA Synthesis Kit (Bio-Rad), according to the manufacturer’s instructions.

For RT-qPCR, samples were prepared using TaqMan™ Fast Advanced Master Mix (Applied Biosystems™) and TaqMan™ probe sets (Applied Biosystems™), according to the manufacturer’s instructions, using the genes summarized in Table 1. Amplifications were carried out in a CFX384 Touch™ Real-Time PCR Detection System (Bio-Rad) using a program of 50°C for 2 min, 95°C for 30 s, and then 40 cycles of 95°C for 3 s and 60°C for 30 s. Each PCR reaction used 2 ng of cDNA. All samples were run in triplicate. No reverse transcriptase and no template controls were included. Threshold cycles were compared to a standard curve generated from serial dilutions of pooled samples for relative quantitation. The relative quantities were normalized to the geometric mean of IPO8, RPL13, and UBC expression.

**Table 1:**
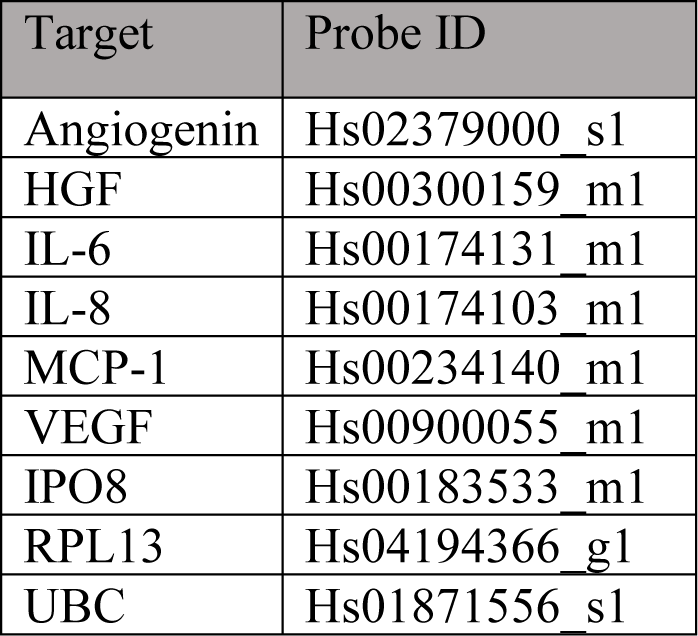
TaqMan™ probes used to assess pro-angiogenic and immunomodulatory gene expression *in vitro*.

| Target | Probe ID |
| --- | --- |
| Angiogenin | Hs02379000_s1 |
| HGF | Hs00300159_m1 |
| IL-6 | Hs00174131_m1 |
| IL-8 | Hs00174103_m1 |
| MCP-1 | Hs00234140_m1 |
| VEGF | Hs00900055_m1 |
| IPO8 | Hs00183533_m1 |
| RPL13 | Hs04194366_g1 |
| UBC | Hs01871556_s1 |

### 2.9 Femoral artery ligation-induced critical limb ischemia (FAL-CLI) model in athymic nude (nu/nu) mice

All animal procedures were performed in accordance with the rules and regulations set by the Canadian Council on Animal Care and were approved by the Animal Care Committee at Western University (Animal Use Protocol #2019-024). *In vivo* assessment of hASC retention and vascular regeneration were performed using 8-week-old athymic *nu/nu* mice (Crl:NU-Foxn1nu, Charles River). To induce unilateral hindlimb ischemia in the mice, femoral artery and vein ligation and excision (2-5 mm) were performed in the right hindlimb, following established protocols [36], [37]. Anesthesia was induced and maintained using isoflurane.

The mice were allowed to recover from the FAL surgery for 24 h, before treatment was applied via intramuscular injection under anesthesia. Six treatment groups were assessed: (i) hASCs in MHA hydrogels, (ii) hASCs in MGC hydrogels, (iii) hASCs in saline, (iv) MHA hydrogel alone controls, (v) MGC hydrogel alone controls, and (vi) saline alone controls. For all treatment groups, a total volume of 20 μL was injected intramuscularly into the surgical limb, just distal to the site of ligation, under anesthesia. Polymer solutions for the hydrogels were prepared as described above for *in vitro* encapsulation. For all treatment groups including hASCs, transduced FLuc^+^ tdT^+^ hASCs were delivered at a concentration of 1.2×10^7^ cells/mL (2.4×10^5^ cells/injection). A total of 29 mice were assessed, with n=5-6 mice/hASC treatment group and n=4 mice/control treatment group.

### 2.10 Laser doppler perfusion imaging assessment of limb perfusion

Longitudinal laser Doppler perfusion imaging (LDPI) was performed to assess hindlimb perfusion at days 1 (prior to treatment), 3, 7, 14, 21, and 28. Mice were anesthetized with isoflurane and warmed at 37°C on a heating pad for 5 min. The mice, maintained under isoflurane anesthesia, were imaged in a supine position using a Moor LDI2-HIR laser Doppler imaging system (Moor Instruments). The average flux in the ischemic (surgical) and control limbs were quantified within a region of interest (ROI) encompassing the foot and ankle joint. The perfusion ratio (PR) was calculated as the ratio between the average flux in the surgical and control limbs and normalized to the perfusion ratio at day 1. All mice included in the study had a day 1 PR <0.15.

### 2.11 In vivo bioluminescence imaging assessment of viable hASC retention

Longitudinal *in vivo* bioluminescence imaging (BLI) was used to assess the retention of the viable FLuc^+^ transduced hASCs in the ischemic limbs at days 1 (following injection), 3, 7, 14, 21, and 28. Mice were anaesthetized using isoflurane and received an intraperitoneal injection of 100 μL of 30 mg/mL D-luciferin (Syd Labs), following which they were imaged using an IVIS Lumina XRMS *In Vivo* Imaging System (PerkinElmer). ROIs of consistent size were drawn around the site of injection and used to quantify the average radiance (p/s/cm^2^/sr) of the luminescence signal. Images were taken using auto-exposure until the average radiance reached a peak value. Average radiance values were normalized to the day 1 values for each mouse.

### 2.12 Immunohistochemical analysis of endothelial cell recruitment

The mice were euthanized 28 days following treatment. The skin covering the hindlimbs was removed, and the entire thigh muscle was harvested and fixed for 24 h in 10% neutral buffered formalin solution (Fisher Scientific). Samples were then subjected to a sucrose gradient (10%, 20%, 30% sucrose in PBS; incubated for 24 h in each solution) before being embedded in frozen section compound (VWR), frozen on dry ice, and stored at −80°C until further processing.

Samples were cryo-sectioned at a thickness of 10 μm and stained to assess host endothelial and mural cell recruitment with antibodies for murine CD31 and α-smooth muscle actin (α-SMA). Sections were incubated in a solution of 1% sodium dodecyl sulphate for 5 min at room temperature, followed by 3 X 5 min washes in PBS. Samples were then blocked in a solution of PBS with 10% donkey serum for 2 h at room temperature. Co-staining for CD31 (1:50 in blocking solution, goat polyclonal anti-CD31, Bio-Techne®, AF3628) and α-SMA (1:100 in blocking solution, rabbit polyclonal anti-αSMA, Abcam AB5694) was performed overnight at 4°C. Sections were washed 3 X 5 min in PBS, and then incubated with the secondary antibodies donkey anti-goat IgG Alexa Fluor™ 647 (1:500 in PBS blocking solution, A-21447, Invitrogen) and donkey anti-rabbit IgG Alexa Fluor™ 488 (1:400 in PBS with 0.05% tween, A-21206, Invitrogen) for 2 h at room temperature. Sections were washed 3 X 5 min in PBS, then mounted using mounting medium with DAPI (Abcam) to stain cell nuclei. Stained cross-sections at 3 depths at least 50 μm apart were imaged for each mouse using an EVOS® FL Imaging microscope (Life Technologies) with a 4X objective. No primary antibody controls were included to confirm staining specificity.

CD31 expression was quantitatively assessed through positive pixel counting within each muscle cross-section using ImageJ. Results were reported as percentage of the positive signal area in the surgical limb, normalized relative to the control (non-surgical limb). Semi-quantitative scoring of the CD31 and α-SMA staining was also performed, to identify and assess vessels observed within the muscle tissue, or in the region of the femoral artery ligation. All scoring was performed in a blinded manner. The tissue sections were assessed based on the size and frequency of vessels showing both CD31 and α-SMA staining, and given scores of 0 (no observed vessels), 1 (low frequency of small vessels), 2 (high frequency of small vessels and/or presence of medium-sized vessels), or 3 (presence of large vessels).

### 2.13 Statistical Methods

All data was presented as the group mean ± standard deviation, unless otherwise noted. For the *in vivo* BLI, individual data points were presented with a line representing the group mean. All statistical analyses were performed using GraphPad Prism 9. Area under the curve (AUC) analysis was performed on the normalized *in vivo* BLI data [38]. Differences between hydrogel groups for the physical comparison assays, and differences between treatment groups for the AUC analysis were assessed using one-way ANOVA with Tukey’s post-hoc test for multiple comparisons. Differences between hydrogel groups at each timepoint for the *in vitro* viability, density, and cytokine release data were analyzed using two-way repeated measures (RM) ANOVA, with Tukey’s post-hoc test for multiple comparisons. Matching was performed across data from the same hASC donors. Differences between treatment groups at each timepoint for the *in vivo* perfusion ratio and BLI average radiance data were detected using two-way RM-ANOVA, with Tukey’s post hoc test for multiple comparisons. Data was considered statistically significant when p < 0.05.

## 3 Results

### 3.1 Hydrogel Characterization

Hydrogels with similar Young’s moduli were prepared to control for potential effects of material stiffness and more specifically assess the impact of polysaccharide hydrogel composition on the encapsulated cells and *in vivo* response. More specifically, formulations of 7.5% (w/v) MHA and 3% (w/v) MGC generated soft hydrogels with similar compressive moduli to the previously-developed MGC+PEG(PTMC-A)_2_ hydrogels (Figure 1A), in the range of ∼4 kPa. In addition, the swelling ratio of all three hydrogel formulations was assessed following overnight incubation in PBS at 37 °C. The MHA hydrogels demonstrated a greater degree of swelling compared to either of the MGC-based hydrogels (Figure 1B), which can be attributed to increased osmotic pressure within the anionic MHA hydrogels relative to the cationic MGC-based formulations [39].

**Figure 1:**
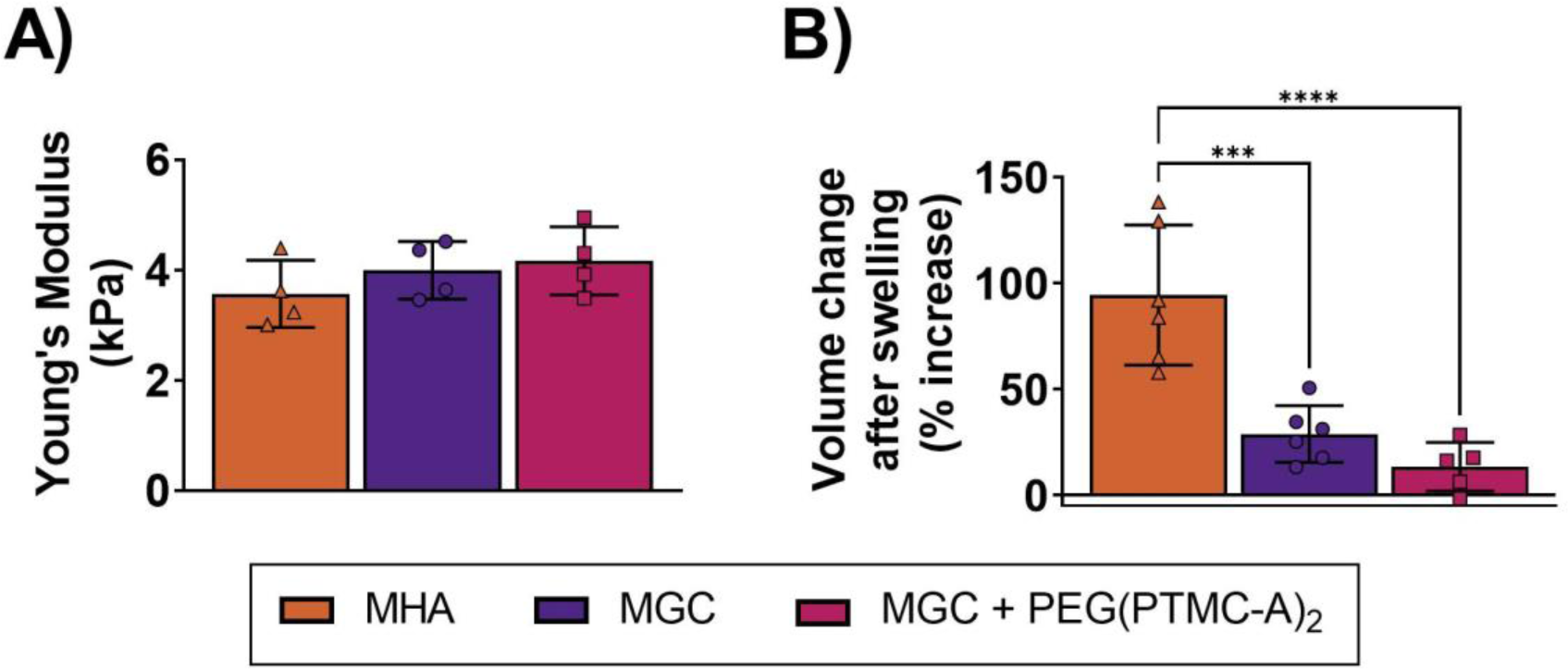
Hydrogels prepared from the three different formulations (7.5% MHA, 3% MGC and 1% MGC + 4% PEG(PTMC-A)_2_) had similar compressive Young’s moduli, but showed different degrees of swelling. **(A)** The average Young’s moduli of the hydrogels determined through compression testing using a CellScale MicroTester. All three hydrogel formulations resulted in soft hydrogels (N=4 hydrogels/formulation, no significant differences between groups). (B) Results from the swelling analysis show that MHA hydrogels swelled to a significantly higher degree than either of the MGC-based hydrogel formulation (N=5-6 hydrogels/formulation). All data presented as mean ± SD. Differences between hydrogel groups were determined using one-way ANOVA, with Tukey’s post-hoc test for multiple comparisons; Significant differences are indicated (*** p<0.001, **** p<0.0001).

### 3.2 Effects of Hydrogel Composition on the Viability of Encapsulated hASCs

To assess the effects of hydrogel composition on cell viability, hASCs were encapsulated in all three hydrogel formulations and cultured for up to 14 days. Representative stitched images following Live/Dead™ staining showed that viable hASCs were homogenously distributed throughout both the MHA and MGC single-phase hydrogels, while hASCs encapsulated in the MGC+PEG(PTMC-A)_2_ hydrogels showed a more heterogenous distribution, likely attributed to phase separation of the materials in the composite hydrogels (Figure 2A). Quantification of cell viability confirmed that the hASCs remained highly viable (>85%) in the MHA and MGC hydrogels for up to 14 days in culture (Figure 2B). hASC viability in the MGC+PEG(PTMC-A)_2_ hydrogels was reduced compared to the single-phase hydrogels at all time points; however an increase in viability from 24 h to days 7 and 14 was observed, likely due to a loss of dead cells over time.

**Figure 2:**
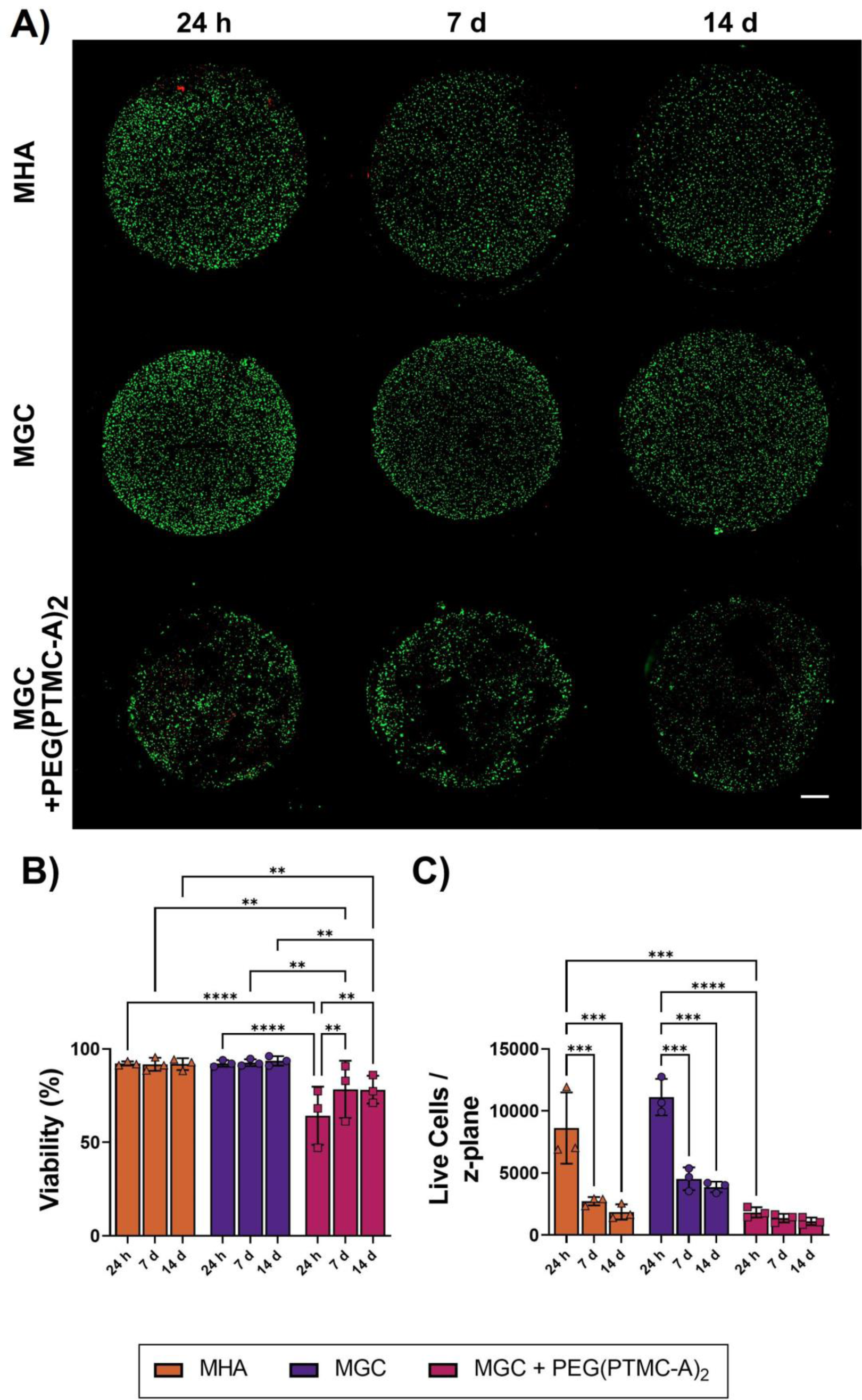
hASCs showed a more homogeneous distribution and higher initial live cell density in the single-phase MGC and MHA hydrogels, with the density declining over time in culture. (A) Live/Dead™ staining was used to visualize hASCs within the three hydrogel formulations across the 14-day culture period. Confocal images (5X) were taken across the entire cross-section in the mid-region of the hydrogels and stitched. Representative images showing viable calcein^+^ cells in green, and dead EthD1^+^ cells in red. Scale bar represents 500 μm. (B) Cell viability was quantified in the stitched images using ImageJ. hASCs remained highly viable (>85%) over 14 days of culture in the MHA and MGC hydrogels. Cell viability was reduced in the MGC+PEG(PTMC-A_2_)_2_ hydrogels compared to the MGC and MHA hydrogels at all time points. (C) Live cell density across the z-plane tiled image was quantified. Both the MHA and MGC hydrogels showed significantly decreased cell densities at 7 and 14 d compared to 24 h. In addition, the hASC density at 24 h was significantly lower in the MGC+PEG(PTMC-A_2_)_2_ hydrogels than in the MGC or MHA hydrogels. All data presented as mean ± SD. Differences between hydrogel groups and timepoints were determined using two-way RM-ANOVA, with Tukey’s correction for multiple comparisons; Significant differences are indicated (** p<0.01, *** p<0.001, **** p<0.0001, n=3-5 replicate hydrogels per timepoint/trial, N=3 trials with different hASC donors).

Live cell density in the hydrogels was calculated based on the total number of live cells across the z-plane at each timepoint. Both single-phase MHA and MGC hydrogels demonstrated decreased live cell densities at days 7 and 14 compared to 24 hours post-encapsulation (Figure 2C). Based on observations of hASCs adhering to the bottom of the well plates, this decline may be attributed at least in part to live cells migrating out of the hydrogels, rather than cell death. At 24 hours post-encapsulation, the live cell density was significantly lower in the composite hydrogels compared to both of the single-phase hydrogels (Figure 2C), which may be related to lower viability or increased cell migration from the composite hydrogels due to the incorporation of the PEG(PTMC-A_2_)_2_ phase.

### 3.3 Effects of Hydrogel Composition on Paracrine Factor Secretion by Encapsulated hASCs

To assess the effects of hydrogel composition on paracrine factor secretion by the encapsulated hASCs, customized Luminex assays and ELISA were used to quantify the levels of paracrine factors associated with pro-angiogenic and immunomodulatory function in CdM samples collected after 48 hours or 7 days of culture (Figure 3). In general, higher levels of paracrine factors were measured in the CdM following 7 days of culture from hASCs encapsulated in both of the MGC-based hydrogels, compared to the CdM from hASCs encapsulated in the MHA hydrogels. At 48 h, significantly more VEGF was released from both MGC-based hydrogel groups compared to the MHA hydrogels, with an even greater amount released from the MGC+PEG(PTMC-A)_2_ hydrogels compared to the single-phase hydrogels. At 7 days, there was significantly higher release of HGF, VEGF, IL-6, and IL-8 from both MGC-based hydrogel formulations compared to the MHA hydrogel group. Additionally at 7 days, there was significantly higher release of angiogenin from the MGC+PEG(PTMC-A_2_)_2_ hydrogels compared to the MHA hydrogels, and significantly higher release of MCP-1 from the single-phase MGC hydrogels compared to the MHA hydrogels.

**Figure 3:**
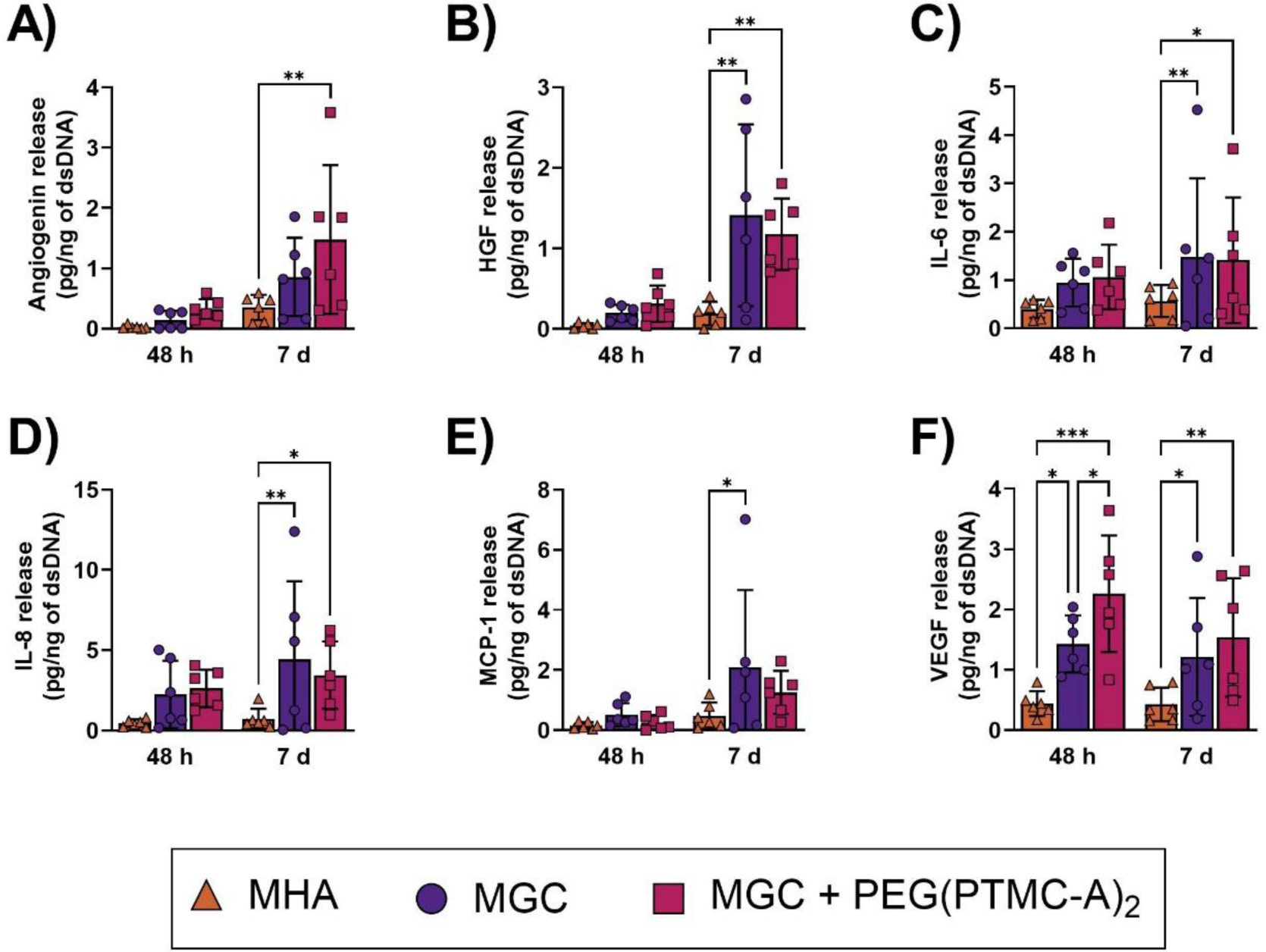
Hydrogel composition influences the levels of pro-angiogenic and immunomodulatory factors detected in conditioned media generated by the encapsulated hASCs. Secreted angiogenin (A), HGF (B), IL-6 (C), IL-8 (D), MCP-1 (E), VEGF (F), in the CdM generated by hASCs encapsulated in the three hydrogel formulations and cultured *in vitro* for 48 h or 7 days, normalized to total dsDNA content. All data presented as mean ± SD. Differences between hydrogel groups at each timepoint were determined using two-way RM-ANOVA, with Tukey’s correction for multiple comparisons; Significant differences are indicated (* p<0.05, ** p<0.01, *** p<0.001, n=3 replicate hydrogels per timepoint/trial, N= 6 trials with different hASC donors).

A small follow-up gene expression study was performed to probe whether the differences measured in the levels of the various secreted factors may be due to changes in the phenotype of the hASCs induced by encapsulation and culture within the different hydrogels or attributed to variable sequestration of the factors within the cationic MGC-based formulations versus anionic MHA hydrogels. Overall, the findings supported that there were differences in phenotype in the hASCs encapsulated within the different hydrogel formulations (Supplementary Figure S2). More specifically, expression of HGF, IL-8, and VEGF were significantly increased in hASCs encapsulated in the MGC hydrogels compared to the MHA hydrogels, consistent with the results from the paracrine factor secretion analysis. The results also support that hydrogel encapsulation alters the hASC pro-angiogenic phenotype compared to 2D culture on TCPS, with significantly reduced expression of IL-6 and MCP-1 following culture in all three hydrogel formulations compared to culture on TCPS, along with significantly enhanced expression of HGF and IL-8.

### 3.4 Characterization of hASC delivery in the MGC and MHA hydrogels in the FAL-CLI model in athymic nude (nu/nu) mice

Following *in vitro* characterization, the FAL-CLI model in athymic nude mice was applied to compare the effects of hydrogel composition on viable hASC retention and pro-angiogenic function *in vivo.* Due to an adverse inflammatory response previously observed in this model when the hASCs were delivered in the composite MGC-RGD+PEG(PTMC-A)_2_ hydrogels [30], only the single-phase MHA and MGC hydrogels were included in the *in vivo* testing, and compared to saline delivery. Mice underwent unilateral FAL surgery, then 24 hours later received a 20 μL injection containing 2.4×10^5^ hASCs delivered in the MHA or MGC hydrogels or saline, or cell-free controls of MHA or MGC hydrogels or saline. Interestingly, the BLI data indicated that hASCs delivered using the MHA hydrogels were better retained than hASCs delivered using the MGC hydrogels or in saline (Figure 4A,B). At day 28, the average cell retention was ∼24% for the hASCs delivered in the MHA hydrogels, as compared to ∼10% and ∼11% for the hASCs delivered in the MGC hydrogels and in saline, respectively. Further, AUC analysis supported there was significantly higher viable hASC retention over the 28-day period in the MHA hydrogel group (Figure 4C).

**Figure 4:**
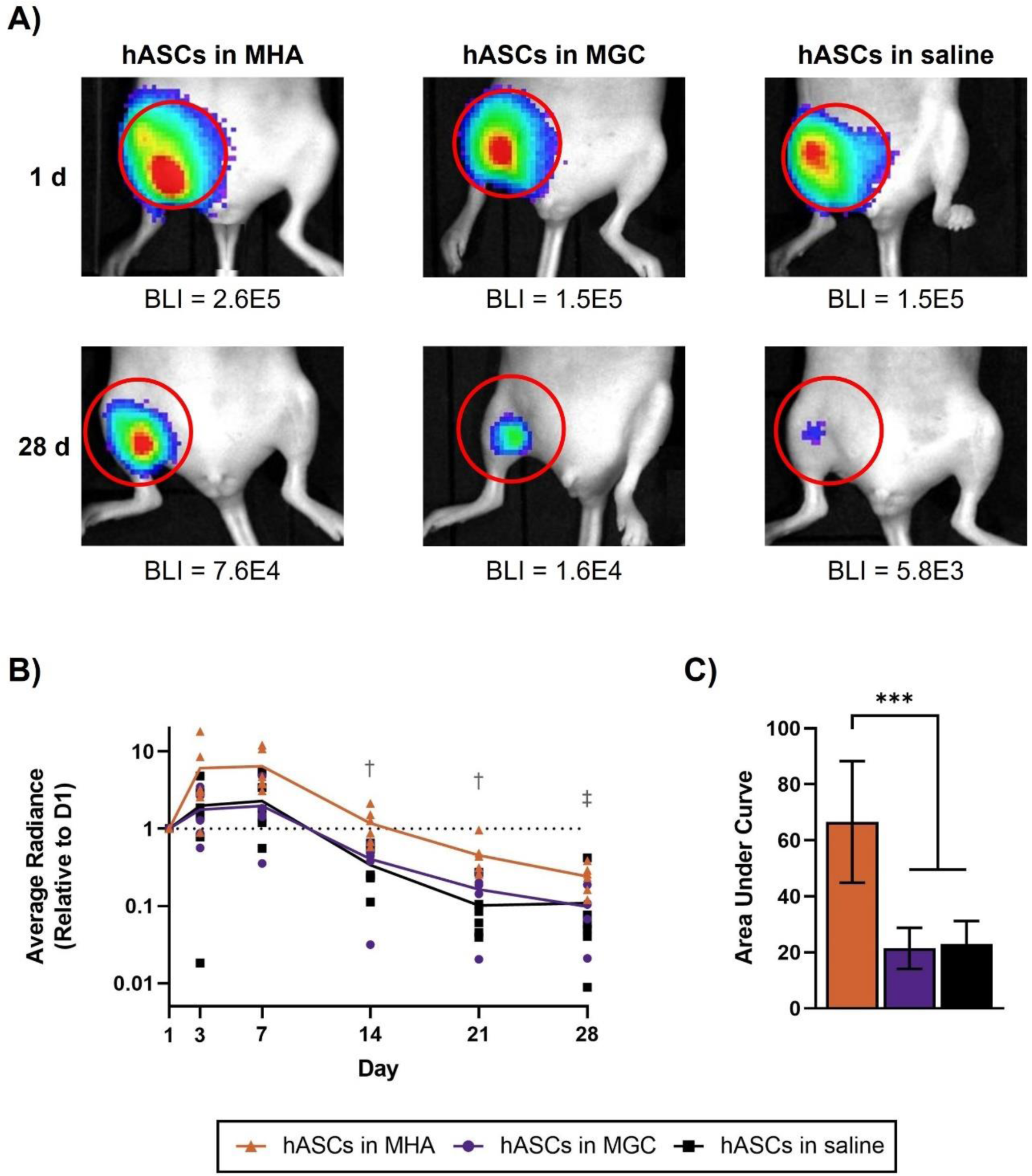
Viable hASCs were better retained following delivery in the MHA hydrogels as compared to delivery in the MGC hydrogels or saline in the FAL-CLI model in *nu/nu* mice. (A) Representative bioluminescent images (BLI) showing FLuc^+^ hASC signal in mice that received hASCs delivered using the MHA or MGC hydrogels, or saline at days 1 and 28. (B) Average bioluminescent radiance (p/s/cm^2^/sr) was calculated within an ROI around the injection site for each mouse at each timepoint, and normalized to values at day 1 (represented by dotted line). hASCs were significantly better retained following delivery in MHA, compared to delivery in saline at days 14 and 21 († p<0.05), and compared to delivery in MGC at day 28 (‡ p<0.05). Individual data points are presented with a line representing the mean. Differences between treatment groups at each timepoint were detected using two-way RM-ANOVA, with Tukey’s post-hoc test for multiple comparisons (n=5-6 mice/hASC treatment group. (C) Area under the curve (AUC) was calculated using the normalized BLI data. Data presented as mean ± SD. Differences between treatment groups were detected using one-way ANOVA, with Tukey’s post-hoc test for multiple comparisons (n=5-6 mice/hASC treatment group).

However, the enhanced cell retention in the MHA group did not translate into enhanced recovery of perfusion. LDPI analysis of hindlimb perfusion over 28 days, quantified using an ROI consisting of the foot region, showed no significant differences between any of the treatment groups at any of the timepoints (Figure 5B).

**Figure 5:**
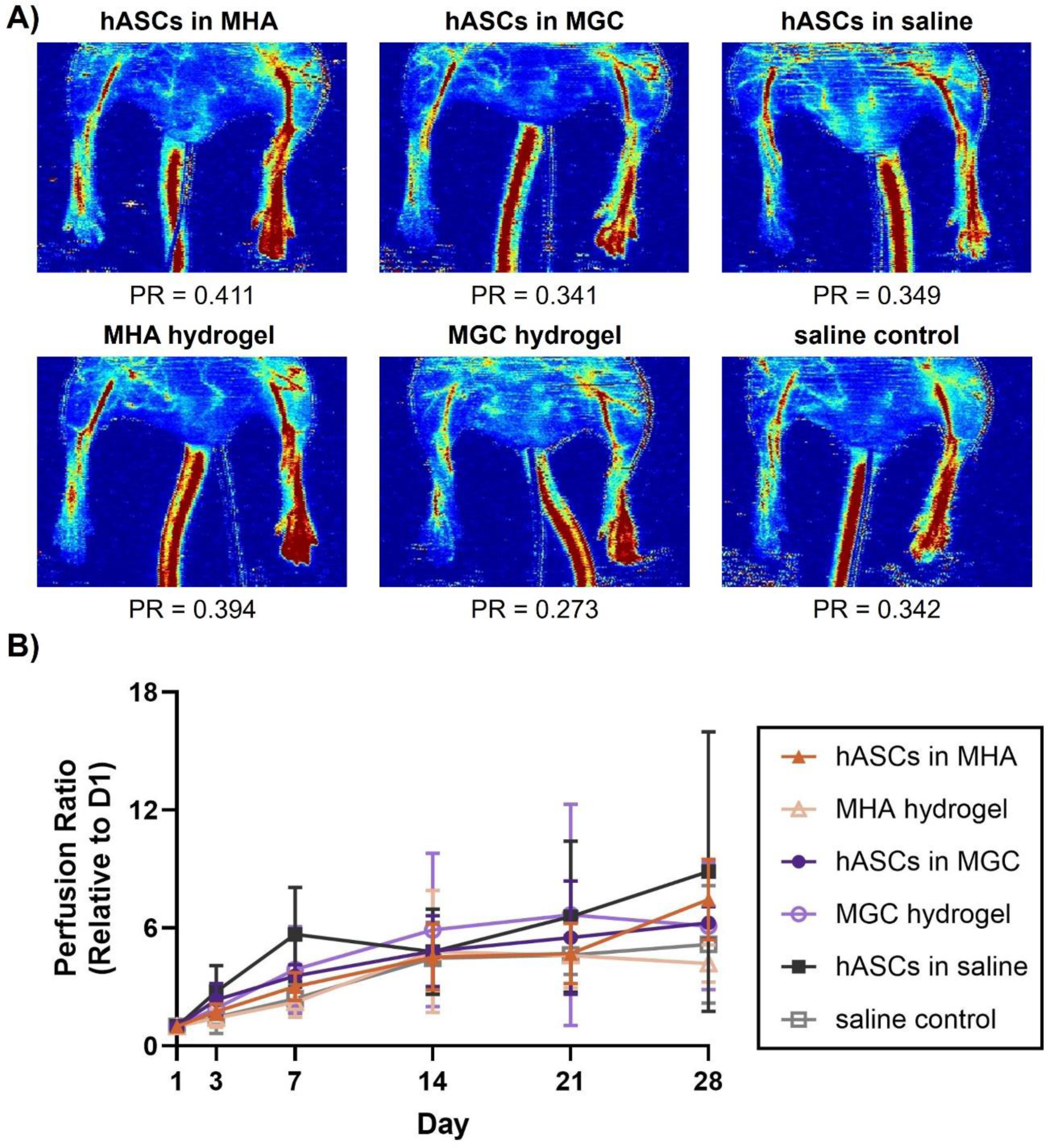
hASC delivery did not accelerate the recovery of limb perfusion in the FAL-CLI model in *nu/nu* mice. (A) Representative LDPI images showing blood flow in the surgical and control limbs at day 28 for each treatment group, along with perfusion ratio (PR) quantifications. (B) Mean flux was quantified within an ROI comprising the foot and ankle for both limbs. PR was calculated from the mean flux of the surgical limb / control limb for each mouse at each time point, and normalized to the perfusion ratio at day 1. No significant differences were observed between the treatment groups at any timepoint. Data presented as mean ± SD. Data was analyzed by two-way RM-ANOVA, with Tukey’s post-hoc test for multiple comparisons (n=5-6 mice/hASC treatment group, n=4 mice/control treatment group).

Immunofluorescence (IF) staining for murine CD31 and α-SMA was performed on cross-sections of the gastrocnemius muscles collected from both the surgical (ischemic) and non-ligated control limbs at day 28. The percent area of CD31^+^ staining was quantified, and the levels in the ligated limbs were normalized to the corresponding control limbs (Figure 6A-B). In general, there was substantial inter-mouse variability, suggesting there was regional variation in the response. Nonetheless, the mean values indicated there were similar levels of CD31^+^ staining in the ischemic and control limbs for mice in all treatment groups.

**Figure 6:**
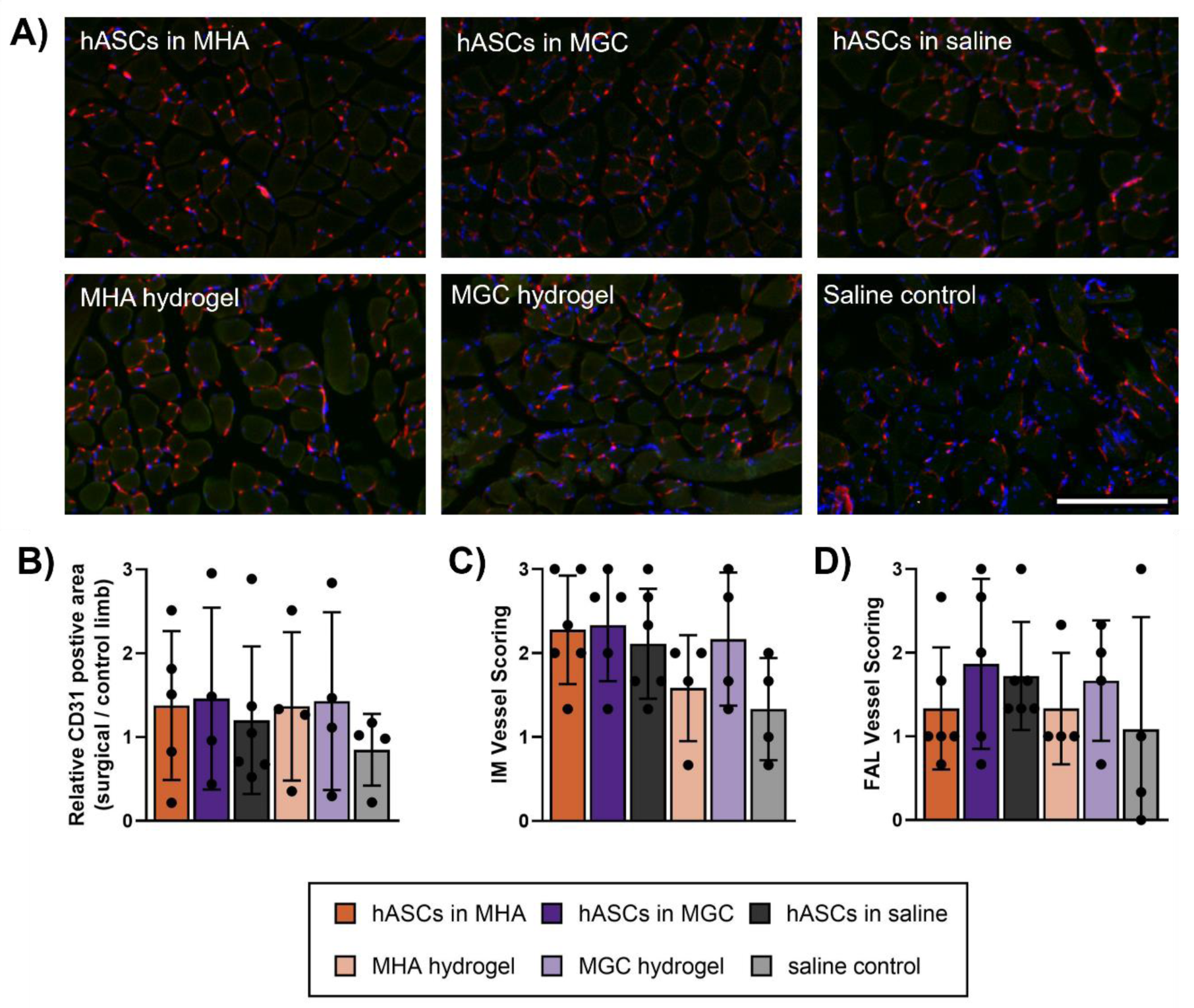
Analysis of CD31 staining suggested that the overall vascular content within ischemic thigh muscles was similar between all treatment groups at day 28. (A) Representative images of thigh muscles: CD31^+^ endothelial cells (red), α-SMA^+^ cells (green), DAPI^+^ cell nuclei (blue). Scale bar represents 200 μm. (B) CD31^+^ area per section, normalized to control limb. (C) Qualitative assessment of frequency and size of intramuscular (IM) vessels was performed and scored on a scale of 0-3, with scores for the triplicate sections averaged for each mouse. (D) Semi-quantitative assessment of the frequency and size of vessels within the femoral artery ligation (FAL) region was performed and scored on a scale of 0-3, with scores for the triplicate sections averaged for each mouse. Individual data points are plotted, with a bar graph representing mean ± SD. No significant differences were observed between treatment groups, as determined using one-way ANOVA, with Tukey’s post-hoc test for multiple comparisons (n=4-6 mice/ treatment group).

In addition, the size and frequency of vessels within the gastrocnemius muscles from the surgical limb (Figure 6C), or within the region of the femoral artery ligation (Figure 6D) were qualitatively assessed. Representative images of vessels of different sizes within the muscle tissue are shown in Supplementary Figure S3. Scoring of vessels showing both CD31 and α-SMA staining was performed in a blinded manner, based on the following scale: 0 (no observed vessels), 1 (low frequency of small vessels), 2 (high frequency of small vessels and/or presence of medium-sized vessels), or 3 (presence of large vessels). While there were no significant differences between the average scores for the treatment groups, no large vessels were observed within the gastrocnemius muscles from the surgical limb of the mice treated with the MHA hydrogel alone or saline alone, suggesting that vascular regeneration may be less advanced in these groups.

## 4 Discussion

Cell delivery within polysaccharide hydrogels is a promising strategy to augment the localized retention of viable cells and help direct their function following intramuscular delivery to ischemic tissues. In designing a hydrogel delivery system for MSCs in the context of PAD/CLI, it is important to consider how the hydrogel material will influence the paracrine function of the MSCs, and how this will further affect the *in vivo* response. Building from our previous work investigating composite MGC-RGD+PEG(PTMC-A)_2_ hydrogels for hASC delivery, this study focused on characterizing the effects of hydrogel composition by comparing the response of hASCs encapsulated within MGC- and MHA-based hydrogels both *in vitro* and *in vivo*. Importantly, the hydrogels were designed to have comparable Young’s moduli to minimize confounding effects of varied material stiffness, which has been shown to influence the MSC secretome [40], [41].

Each of the platforms supported hASC viability following encapsulation and *in vitro* culture over 14 days. Notably, the cells were more heterogeneously distributed and had significantly lower viability in the composite MGC+PEG(PTMC-A)_2_ hydrogels as compared to the single-phase MGC and MHA hydrogels. Similarly, Anjum *et al*. showed that the incorporation of PEG(PTMC-A)_2_ into methacrylated chondroitin sulphate (MCS) hydrogels reduced the viability of encapsulated bovine ASCs [29], suggesting that the PEG(PTMC-A)_2_ may negatively influence ASC survival. Analysis of live cell density showed a significant reduction in the MGC and MHA groups from 24 h to 7 days, with the levels then sustained out to 14 days. The identification of cells seeding the well floor suggested that some of the cells were migrating out of the gels. Despite this decline in cell density, we proceeded with functional characterization experiments *in vitro*, as a high density of viable cells was maintained within the hydrogels over time. Moreover, it remains unclear how long the cells should persist *in vivo* to maximize their therapeutic effects, and it is possible that cell migration out of the hydrogels could be beneficial for *in vivo* regeneration.

There has been very limited research to date on how hydrogel composition influences paracrine factor expression in encapsulated cell populations. Most studies assess only a single type of hydrogel, and comparison between studies is difficult due to differences in multiple factors including cell type and density, material composition and functionalization, specific methods of assessment, and timepoints, all of which can influence the observed response. The enhanced expression of pro-angiogenic and immunomodulatory factors detected in the CdM samples from the MGC-based materials relative to the MHA hydrogels suggests that the MGC may be having an activating or priming effect on the hASCs. Interestingly, chitosan and chitin have been investigated as inflammatory priming stimuli for MSCs [42], [43]. More specifically, Farrokhi *et al*. showed that murine ASCs cultured on chitosan films *in vitro* secreted significantly more TGF-β and IL-10, and had higher arginase activity compared to MSCs that were primed with lipopolysaccharide or untreated [42]. In another study, Lieder *et al*. showed that *in vitro* stimulation with chitin and chitosan oligosaccharides enhanced the secretion of IL-6 and IL-8 from human bone marrow-derived (BM) MSCs compared to unstimulated controls [43]. It is important to note that the levels of paracrine factors detected in the CdM samples in the current study could be impacted by differences in what is being released from the hydrogels into the media, rather than differences in hASC secretion. More specifically, the anionic HA hydrogels would be more likely to sequester positively-charged growth factors, as compared to the cationic MGC-based hydrogels. However, the follow-up gene expression study supported that there were phenotypic differences between the hASCs encapsulated and cultured in the different hydrogel formulations, with enhanced expression of HGF, VEGF, and IL-8 in the MGC hydrogels compared to the MHA hydrogels.

While there is growing interest in the development of strategies to augment paracrine factor secretion by MSC populations [44], it remains unclear whether the increased secretion of various factors will ultimately augment functional tissue regeneration. Notably, many of the growth factors and cytokines investigated can have varied effects on different cell populations. The combined effects of these factors could potentially be beneficial, with outcomes likely dependent on concentrations, as well as timing of release. In particular, there is a close interrelationship between angiogenesis and inflammation. While chronic inflammation can have negative effects on vascular structure and function [45], acute inflammation has been shown to play an important role in initiating angiogenesis [46]. As such, the secretion of factors such as IL-6, IL-8, and MCP-1 that can have pro-inflammatory effects, including promoting the recruitment of immune cell populations such as neutrophils and macrophages [47], may also help to stimulate vascular regeneration. In addition, IL-6 and HGF have been shown to modulate macrophage polarization and have the capacity to promote a more pro-regenerative M2-like macrophage phenotype [48], [49]. However, there is a fine balance with the potential for cascading effects, and the sustained presence of high levels of pro-inflammatory factors could ultimately be detrimental for the formation of mature functional vessels [50], [51].

Interestingly, the BLI data from the *in vivo* study indicated that hASC retention was augmented when the cells were delivered in the MHA hydrogels as compared to delivery in the MGC hydrogels or in saline. The improved cell retention in the MHA hydrogels relative to saline suggests that the MHA was having protective effects, supporting the localized survival of the hASCs within the FAL-CLI model. Other studies have shown that hydrogels can improve the retention of delivered MSCs, as measured by BLI or longitudinal fluorescence cell tracking. For example, Huang *et al*. showed using BLI that self-assembled peptide hydrogels enhanced the retention of human placenta-derived MSCs delivered to an FAL-CLI model in BALB/c nude mice in comparison to delivery in saline [15]. Similarly, Wang *et al*. showed using longitudinal fluorescence imaging that hydrogels composed of chondroitin sulphate and hyaluronic acid enhanced the retention of mouse ASCs following delivery to an FAL-CLI model in BALB/c nude mice relative to saline delivery controls [12].

In contrast to the MHA, the MGC did not improve the localized retention of viable hASCs within our *in vivo* model. It is possible that the enhanced secretion levels of the pro-inflammatory factors from the MGC hydrogels as compared to the MHA hydrogels may have influenced this response. In particular, the augmented secretion of factors such as MCP-1 and IL-6 may have stimulated the recruitment and activation of endogenous macrophages, accelerating the clearance of the delivered hASCs. This interpretation is supported by our previous work, which showed enhanced recruitment and greater infiltration of CD68^+^ macrophages following delivery of rat ASCs within the composite MGC-RGD+PEG(PTMC-A)_2_ hydrogels in a healthy immunocompetent rat model, as compared to hydrogel alone controls [23]. Moreover, it has been suggested that differences in the overall charge of the hydrogels can alter the activity of endogenous macrophages. More specifically, Brodbeck *et al*. demonstrated in an *in vivo* implant model that anionic polymers showed decreased macrophage adhesion and fusion, as well as increased macrophage apoptosis, compared to neutral and cationic polymers [52]. Therefore, the cationic MGC hydrogels may have more strongly promoted macrophage recruitment and activation as compared to the anionic MHA hydrogels, leading to increased hASC clearance due to macrophage phagocytosis.

While the current study focused on analyzing markers of vascular regeneration in tissues collected at the endpoint of 28 days, it would be interesting to explore differences in the host macrophage response between the MGC and MHA hydrogels in future work. For example, Sehl *et al*., demonstrated the feasibility of using multi-modal imaging to detect and quantify delivered MSCs over time, while simultaneously characterizing the infiltration of host macrophages [53]. More specifically, MSCs labelled with iron oxide nanoparticles were detected using magnetic resonance imaging and magnetic particle imaging, while macrophages were labelled using perfluorocarbon and detected using fluorine-19 magnetic resonance imaging [53]. A similar strategy could be applied in the future to assess the correlation between the host macrophage response and hASC retention in the different hydrogel formulations. Moreover, IF staining could be performed to further characterize the macrophage response, using the murine macrophage marker F4/80 and the phagocytic macrophage marker CD68, in combination with phenotypic markers such as the M1-like markers CD80 and iNOS, and the M2-like markers CD163 and Arg-1 [54], [55].

Notably, longitudinal LDPI imaging and end-point analysis of CD31 staining within the gastrocnemius muscles showed that the enhanced hASC retention in the MHA group did not translate to an enhanced recovery of limb perfusion or changes in the vessel density at 28 days in the current model. There was a reduced presence of larger vessels within the muscles in the surgical limb of the mice that were treated with MHA alone or saline alone, suggesting there may be reduced vascular regeneration in these groups. However, a high degree of inter-animal variability was observed in both the LDPI and IF data. This is consistent with the observations of Zbinden *et al*., who showed using BALB/c mice that there is large inter-mouse variability in the functionality of pre-existing collaterals, and that this can greatly impact the assessment of therapies using the FAL-CLI model [56].

Overall, the results indicate that enhanced cell retention was insufficient to augment vascular regeneration within our model. These findings are in contrast to other studies that have shown that MSC delivery within hydrogels enhances MSC retention and is also associated with increased vascular regeneration in various FAL-CLI models [12], [13], [15], [57]. One possible reason for this discrepancy may be differences in the cell dose that was delivered. For example, in studies reporting positive outcomes, Wang *et al*. delivered 10^6^ murine ASCs per mouse, while Huang *et al*. delivered 3×10^7^ human placenta-derived MSCs per mouse, which are both substantially greater than the cell dose of 2.4×10^5^ hASCs per mouse in the current study [12], [15]. Notably, there have been few studies to date that have systematically characterized the cell dose required to stimulate vascular regeneration. Interestingly, Moon *et al.* delivered 10^5^, 5×10^5^, or 10^6^ hASCs to an FAL-CLI model in athymic nude mice, and showed that limb perfusion increased according to the number of transplanted cells, with effects starting at the lowest cell dose in comparison to saline controls [8]. In general, further research is critically needed to establish targets for a therapeutic cell dose, which may be different if the cells are delivered using a biomaterial versus saline. It is important to recognize that these targets will also depend on the specific MSC source applied, as well as the characteristics of the cell donor, including age, sex, and health status [58]–[60]. Overall, there is a need for the field to establish potency indicators for the specific cell population(s) of interest, to select donor cells that have the functional capacity to reliably and predictably stimulate regeneration and limit issues associated with cell donor variability [61].

## 5 Conclusion

In conclusion, this study demonstrated that hydrogel composition can alter the viability, paracrine function, and *in vivo* retention of encapsulated hASCs. Notably, hydrogel composition was shown to influence paracrine factor secretion, with greater levels of pro-angiogenic and immunomodulatory factors detected in the conditioned media samples from the hASCs encapsulated in the MGC-based hydrogels as compared to the MHA hydrogels. Future studies should incorporate functional cell-based assays to better understand how the altered paracrine secretion levels may influence key endogenous cell populations involved in the host regenerative response to develop a deeper understanding of the underlying mechanisms of hASC-mediated regeneration. For example, the effects of the CdM on human microvascular endothelial cell survival and tubule formation could be assessed to probe pro-angiogenic function. Additionally, the effects of the CdM on the phenotype, migration, and phagocytic activity of human macrophages could be assessed as a measure of immunomodulatory function. Following delivery to an FAL-CLI model in athymic *nu/nu* mice, the hASCs were better retained within the MHA hydrogels as compared to delivery in the MGC hydrogels or saline. However, enhanced cell retention was not associated with augmented recovery of limb perfusion or changes in vessel density, based on LDPI analysis and CD31 staining. One potential reason for the limited regenerative response observed is the relatively low dose of cells delivered in the current study. Further studies are warranted to systematically assess whether higher initial cell doses and/or repeated treatments would achieve greater therapeutic effects, with consideration given to feasibility in scaling up and translating these approaches for clinical use in humans.

## Data Availability Statement

The raw data supporting the conclusions of this manuscript will be made available by the authors, without undue reservation, to any qualified researcher.

## Acknowledgements

This study was supported by Grant-in-Aid funding from the Heart and Stroke Foundation of Canada (G-19-0026269) and the Canadian Institutes of Health Research (CIHR MOP #378189). Scholarship support for Fiona Serack was provided through an Ontario Graduate Scholarship and a Transdisciplinary Bone & Joint Training Award from the Bone and Joint Institute at Western University. The authors thank Drs. A. Grant and D. Matic for their clinical collaborations with the provision of adipose tissue samples required for cell isolation, as well as Drs. Ying Xia, John Kelly, and Yue Li for their technical support. Additionally, the authors would like to thank Dr. Eric Arts for access to the Bio-Plex® MAGPIX™ Multiplex Reader system for the Luminex® analyses and the London Regional Microscopy Facility for access to laser Doppler perfusion imaging equipment.

## References

[1] P. Song et al., “Global, regional, and national prevalence and risk factors for peripheral artery disease in 2015: an updated systematic review and analysis,” Lancet Glob. Heal., vol. 7, no. 8, pp. e1020–e1030, 2019.

[2] V. N. Varu, M. E. Hogg, and M. R. Kibbe, “Critical limb ischemia,” J. Vasc. Surg., vol. 51, no. 1, pp. 230–241, 2010.

[3] S. Duff, M. S. Mafilios, P. Bhounsule, and J. T. Hasegawa, “The burden of critical limb ischemia: A review of recent literature,” Vasc. Health Risk Manag., vol. 15, pp. 187–208, 2019.

[4] Z. Shirbaghaee, M. Hassani, S. Heidari Keshel, and M. Soleimani, “Emerging roles of mesenchymal stem cell therapy in patients with critical limb ischemia,” Stem Cell Res. Ther., vol. 13, no. 1, p. 462, 2022.

[5] J. M. Gimble, A. J. Katz, and B. A. Bunnell, “Adipose-derived stem cells for regenerative medicine,” Circulation Research, vol. 100, no. 9. pp. 1249–1260, 2007.

[6] Y. Ikegame et al., “Comparison of mesenchymal stem cells from adipose tissue and bone marrow for ischemic stroke therapy,” Cytotherapy, vol. 13, no. 6, pp. 675–685, 2011.

[7] Y. J. Kim, H. K. Kim, H. K. Cho, Y. C. Bae, K. T. Suh, and J. S. Jung, “Direct comparison of human mesenchymal stem cells derived from adipose tissues and bone marrow in mediating neovascularization in response to vascular ischemia,” Cell. Physiol. Biochem., vol. 20, no. 6, pp. 867–876, 2007.

[8] M. H. Moon et al., “Human adipose tissue-derived mesenchymal stem cells improve postnatal neovascularization in a mouse model of hindlimb ischemia,” Cell. Physiol. Biochem., vol. 17, no. 5–6, pp. 279–290, 2006.

[9] W. Fan et al., “Adipose Stromal Cells Amplify Angiogenic Signaling via the VEGF/mTOR/Akt Pathway in a Murine Hindlimb Ischemia Model: A 3D Multimodality Imaging Study,” PLoS One, vol. 7, no. 9, 2012.

[10] M. Qadura, D. C. Terenzi, S. Verma, M. Al-Omran, and D. A. Hess, “Concise Review: Cell Therapy for Critical Limb Ischemia: An Integrated Review of Preclinical and Clinical Studies,” Stem Cells, vol. 36, no. 2, pp. 161–171, 2018.

[11] J. P. Laurila et al., “Human embryonic stem cell-derived mesenchymal stromal cell transplantation in a rat hind limb injury model,” Cytotherapy, vol. 11, no. 6, pp. 726–737, 2009.

[12] X. Wang et al., “Composite Hydrogel Modified by IGF-1C Domain Improves Stem Cell Therapy for Limb Ischemia,” ACS Appl. Mater. Interfaces, vol. 10, no. 5, pp. 4481–4493, 2018.

[13] S. A. Young et al., “Mechanically resilient injectable scaffolds for intramuscular stem cell delivery and cytokine release,” Biomaterials, vol. 159, pp. 146–160, 2018.

[14] Y. Li et al., “Primed 3D injectable microniches enabling low-dosage cell therapy for critical limb ischemia,” Proc. Natl. Acad. Sci. U. S. A., vol. 111, no. 37, pp. 13511–13516, 2014.

[15] A. Huang et al., “Self-assembled GFFYK peptide hydrogel enhances the therapeutic efficacy of mesenchymal stem cells in a mouse hindlimb ischemia model,” Acta Biomater., vol. 85, pp. 94–105, 2019.

[16] S. R. Caliari and J. A. Burdick, “A practical guide to hydrogels for cell culture,” Nat. Methods, vol. 13, no. 5, pp. 405–414, 2016.

[17] Y. Okamoto, M. Watanabe, K. Miyatake, M. Morimoto, Y. Shigemasa, and S. Minami, “Effects of chitin/chitosan and their oligomers/monomers on migrations of fibroblasts and vascular endothelium,” Biomaterials, vol. 23, no. 9, pp. 1975–1979, 2002.

[18] G. Tripodo et al., “Hydrogels for biomedical applications from glycol chitosan and PEG diglycidyl ether exhibit pro-angiogenic and antibacterial activity,” Carbohydr. Polym., vol. 198, pp. 124–130, 2018.

[19] Z. Liu et al., “The influence of chitosan hydrogel on stem cell engraftment, survival and homing in the ischemic myocardial microenvironment,” Biomaterials, vol. 33, no. 11, pp. 3093–3106, 2012.

[20] X. Chen et al., “Peptide SIKVAV-modified chitosan hydrogels promote skin wound healing by accelerating angiogenesis and regulating cytokine secretion,” Am. J. Transl. Res., vol. 10, no. 12, pp. 4258–4268, Dec. 2018.

[21] N. Zhao et al., “IGF-1C domain-modified hydrogel enhances therapeutic potential of mesenchymal stem cells for hindlimb ischemia,” Stem Cell Res. Ther., vol. 10, no. 1, 2019.

[22] W. N. Lu et al., “Functional improvement of infarcted heart by co-injection of embryonic stem cells with temperature-responsive chitosan hydrogel,” Tissue Eng. - Part A, vol. 15, no. 6, pp. 1437–1447, 2009.

[23] S. A. Young, L. E. Flynn, and B. G. Amsden, “ Adipose-Derived Stem Cells in a Resilient In Situ Forming Hydrogel Modulate Macrophage Phenotype,” Tissue Eng. Part A, vol. 24, no. 23–24, pp. 1784–1797, 2018.

[24] K. T. Dicker, L. A. Gurski, S. Pradhan-Bhatt, R. L. Witt, M. C. Farach-Carson, and X. Jia, “Hyaluronan: A simple polysaccharide with diverse biological functions,” Acta Biomater., vol. 10, no. 4, pp. 1558–1570, 2014.

[25] D. C. West and S. Kumar, “Hyaluronan and angiogenesis.,” Novartis Found. Symp., vol. 143, 1989.

[26] J. S. Frenkel, “The role of hyaluronan in wound healing,” Int. Wound J., vol. 11, no. 2, pp. 159–163, 2014.

[27] Y. Li, Y. Xiao, and C. Liu, “The Horizon of Materiobiology: A Perspective on Material-Guided Cell Behaviors and Tissue Engineering,” Chem. Rev., vol. 117, no. 5, pp. 4376–4421, 2017.

[28] E. S. Place, N. D. Evans, and M. M. Stevens, “Complexity in biomaterials for tissue engineering,” Nat. Mater., vol. 8, no. 6, pp. 457–470, 2009.

[29] F. Anjum, A. Carroll, S. A. Young, L. E. Flynn, and B. G. Amsden, “Tough, Semisynthetic Hydrogels for Adipose Derived Stem Cell Delivery for Chondral Defect Repair,” Macromol. Biosci., vol. 17, no. 5, pp. 1–13, 2017.

[30] F. E. Serack, J. A. Ronald, B. G. Amsden, D. A. Hess, and L. E. Flynn, “Delivery of human adipose-derived stromal cells within mechanically resilient hydrogels induces adverse outcomes in a femoral artery ligation model in athymic nu/nu mice,” bioRxiv, 2023.

[31] M. Costantini et al., “3D bioprinting of BM-MSCs-loaded ECM biomimetic hydrogels for in vitro neocartilage formation,” Biofabrication, vol. 8, no. 3, 2016.

[32] C. C. L. Schuurmans, M. Mihajlovic, C. Hiemstra, K. Ito, W. E. Hennink, and T. Vermonden, “Hyaluronic acid and chondroitin sulfate (meth)acrylate-based hydrogels for tissue engineering: Synthesis, characteristics and pre-clinical evaluation,” Biomaterials, vol. 268, 2021.

[33] B. Johnson, J. M. Bauer, D. J. Niedermaier, W. C. Crone, and D. J. Beebe, “Experimental techniques for mechanical characterization of hydrogels at the microscale,” Exp. Mech., vol. 44, no. 1, pp. 21–28, 2004.

[34] L. Flynn, J. L. Semple, and K. A. Woodhouse, “Decellularized placental matrices for adipose tissue engineering,” J. Biomed. Mater. Res. - Part A, vol. 79, no. 2, pp. 359–369, 2006.

[35] A. M. Hamilton, P. J. Foster, and J. A. Ronald, “Evaluating Nonintegrating Lentiviruses as Safe Vectors for Noninvasive Reporter-Based Molecular Imaging of Multipotent Mesenchymal Stem Cells,” Hum. Gene Ther., vol. 29, no. 10, pp. 1213–1225, 2018.

[36] T. Couffinhal, M. Silver, L. P. Zheng, M. Kearney, B. Witzenbichler, and J. M. Isner, “Mouse model of angiogenesis.,” Am. J. Pathol., vol. 152, no. 6, pp. 1667–1679, Jun. 1998.

[37] B. J. Capoccia, R. M. Shepherd, and D. C. Link, “G-CSF and AMD3100 mobilize monocytes into the blood that stimulate angiogenesis in vivo through a paracrine mechanism,” Blood, vol. 108, no. 7, pp. 2438–2445, Oct. 2006.

[38] M. Köberle, K. Müller, M. Kamprad, F. Horn, and M. Scholz, “Monitoring disease progression and therapeutic response in a disseminated tumor model for non-hodgkin lymphoma by bioluminescence imaging,” Mol. Imaging, vol. 14, no. 6, pp. 400–411, 2015.

[39] S. Morozova and M. Muthukumar, “Elasticity at Swelling Equilibrium of Ultrasoft Polyelectrolyte Gels: Comparisons of Theory and Experiments,” Macromolecules, vol. 50, no. 6, pp. 2456–2466, 2017.

[40] G. D. Kusuma, J. Carthew, R. Lim, and J. E. Frith, “Effect of the Microenvironment on Mesenchymal Stem Cell Paracrine Signaling: Opportunities to Engineer the Therapeutic Effect,” Stem Cells Dev., vol. 26, no. 9, pp. 617–631, 2017.

[41] M. E. Wechsler, V. V. Rao, A. N. Borelli, and K. S. Anseth, “Engineering the MSC Secretome: A Hydrogel Focused Approach,” Adv. Healthc. Mater., vol. 10, no. 7, 2021.

[42] S. Farrokhi, F. sotoodehnejadnematalahi, A. Fathollahi, M. Haji Molla Hoseini, S. M. Hashemi, and F. Yeganeh, “The immunomodulatory potential of murine adipose-derived mesenchymal stem cells is enhanced following culture on chitosan film,” Tissue Cell, vol. 74, 2022.

[43] R. Lieder et al., “Chitosan and Chitin Hexamers affect expansion and differentiation of mesenchymal stem cells differently,” Int. J. Biol. Macromol., vol. 51, no. 4, pp. 675–680, 2012.

[44] N. D. C. Noronha et al., “Correction to: Priming approaches to improve the efficacy of mesenchymal stromal cell-based therapies (Stem Cell Research and Therapy (2019) 10 (131) DOI: 10.1186/s13287-019-1224-y),” Stem Cell Res. Ther., vol. 10, no. 1, pp. 1–21, 2019.

[45] X. Castellon and V. Bogdanova, “Chronic inflammatory diseases and endothelial dysfunction,” Aging Dis., vol. 7, no. 1, 2016.

[46] S. Frantz, K. A. Vincent, O. Feron, and R. A. Kelly, “Innate immunity and angiogenesis,” Circ. Res., vol. 96, no. 1, pp. 15–26, 2005.

[47] M. D. Turner, B. Nedjai, T. Hurst, and D. J. Pennington, “Cytokines and chemokines: At the crossroads of cell signalling and inflammatory disease,” Biochim. Biophys. Acta - Mol. Cell Res., vol. 1843, no. 11, pp. 2563–2582, 2014.

[48] J. J. Fuster and K. Walsh, “The Good, the Bad, and the Ugly of interleukin-6 signaling,” EMBO J., vol. 33, no. 13, pp. 1425–1427, 2014.

[49] W. Choi, J. Lee, J. Lee, S. H. Lee, and S. Kim, “Hepatocyte growth factor regulates macrophage transition to the M2 phenotype and promotes murine skeletal muscle regeneration,” Front. Physiol., vol. 10, no. JUL, 2019.

[50] G. K. Hansson, “Inflammation, Atherosclerosis, and Coronary Artery Disease,” N. Engl. J. Med., vol. 352, no. 16, pp. 1685–1695, 2005.

[51] P. Martin and D. B. Gurevich, “Macrophage regulation of angiogenesis in health and disease,” Semin. Cell Dev. Biol., vol. 119, pp. 101–110, 2021.

[52] W. G. Brodbeck et al., “Biomaterial adherent macrophage apoptosis is increased by hydrophilic and anionic substrates in vivo,” Proc. Natl. Acad. Sci. U. S. A., vol. 99, no. 16, pp. 10287–10292, 2002.

[53] O. C. Sehl, A. V. Makela, A. M. Hamilton, and P. J. Foster, “Trimodal cell tracking in vivo: Combining iron-and fluorine-based magnetic resonance imaging with magnetic particle imaging to monitor the delivery of mesenchymal stem cells and the ensuing inflammation,” Tomography, vol. 5, no. 4, pp. 367–376, 2019.

[54] M. L. Novak and T. J. Koh, “Macrophage phenotypes during tissue repair,” J. Leukoc. Biol., vol. 93, no. 6, pp. 875–881, 2013.

[55] T. H. Tran, R. Rastogi, J. Shelke, and M. M. Amiji, “Modulation of Macrophage Functional Polarity towards Anti-Inflammatory Phenotype with Plasmid DNA Delivery in CD44 Targeting Hyaluronic Acid Nanoparticles,” Sci. Rep., vol. 5, 2015.

[56] S. Zbinden et al., “Interanimal variability in preexisting collaterals is a major factor determining outcome in experimental angiogenesis trials,” Am. J. Physiol. - Hear. Circ. Physiol., vol. 292, no. 4, pp. 1891–1897, 2007.

[57] J. Wang et al., “A cellular delivery system fabricated with autologous BMSCs and collagen scaffold enhances angiogenesis and perfusion in ischemic hind limb,” J. Biomed. Mater. Res. - Part A, vol. 100 A, no. 6, pp. 1438–1447, 2012.

[58] J. Galipeau and L. Sensébé, “Mesenchymal Stromal Cells: Clinical Challenges and Therapeutic Opportunities,” Cell Stem Cell, vol. 22, no. 6, pp. 824–833, 2018.

[59] M. Najar et al., “Therapeutic Mesenchymal Stem/Stromal Cells: Value, Challenges and Optimization,” Front. Cell Dev. Biol., vol. 9, 2022.

[60] P. Nammian, S. L. Asadi-Yousefabad, S. Daneshi, M. H. Sheikhha, S. M. B. Tabei, and V. Razban, “Comparative analysis of mouse bone marrow and adipose tissue mesenchymal stem cells for critical limb ischemia cell therapy,” Stem Cell Res. Ther., vol. 12, no. 1, 2021.

[61] K. P. Robb, J. Audet, R. Gandhi, and S. Viswanathan, “Putative critical quality attribute matrix identifies mesenchymal stromal cells with potent immunomodulatory and angiogenic ‘fitness’ ranges in response to culture process parameters,” Front. Immunol., vol. 13, 2022.

