## Supplementary Table S1 for "Probing the effects of polysaccharide hydrogel composition on the viability and pro-angiogenic function of human adipose-derived stromal cells"

Supplementary Table S1. Donor list for all studies

| Donor | Gender | Age | BMI | Explant Site | Experiments |
| --- | --- | --- | --- | --- | --- |
| 1 | Female | 57 | 24.5 | abdomen | - *in vitro* viability - *in vitro* secretome - *in vivo* study |
| 2 | Female | 65 | 26.2 | breast | - *in vitro* viability - *in vitro* secretome - *in vivo* study |
| 3 | Female | 43 | 41 | abdomen | - *in vitro* viability - *in vitro* secretome |
| 4 | Female | 26 | 26.8 | abdomen | - *in vitro* secretome - *in vivo* study |
| 5 | Female | 51 | 28.5 | abdomen | - *in vitro* secretome |
| 6 | Female | 49 | 27.5 | breast | - *in vitro* secretome |
