## Supplementary Figure S1 for "Probing the effects of polysaccharide hydrogel composition on the viability and pro-angiogenic function of human adipose-derived stromal cells"

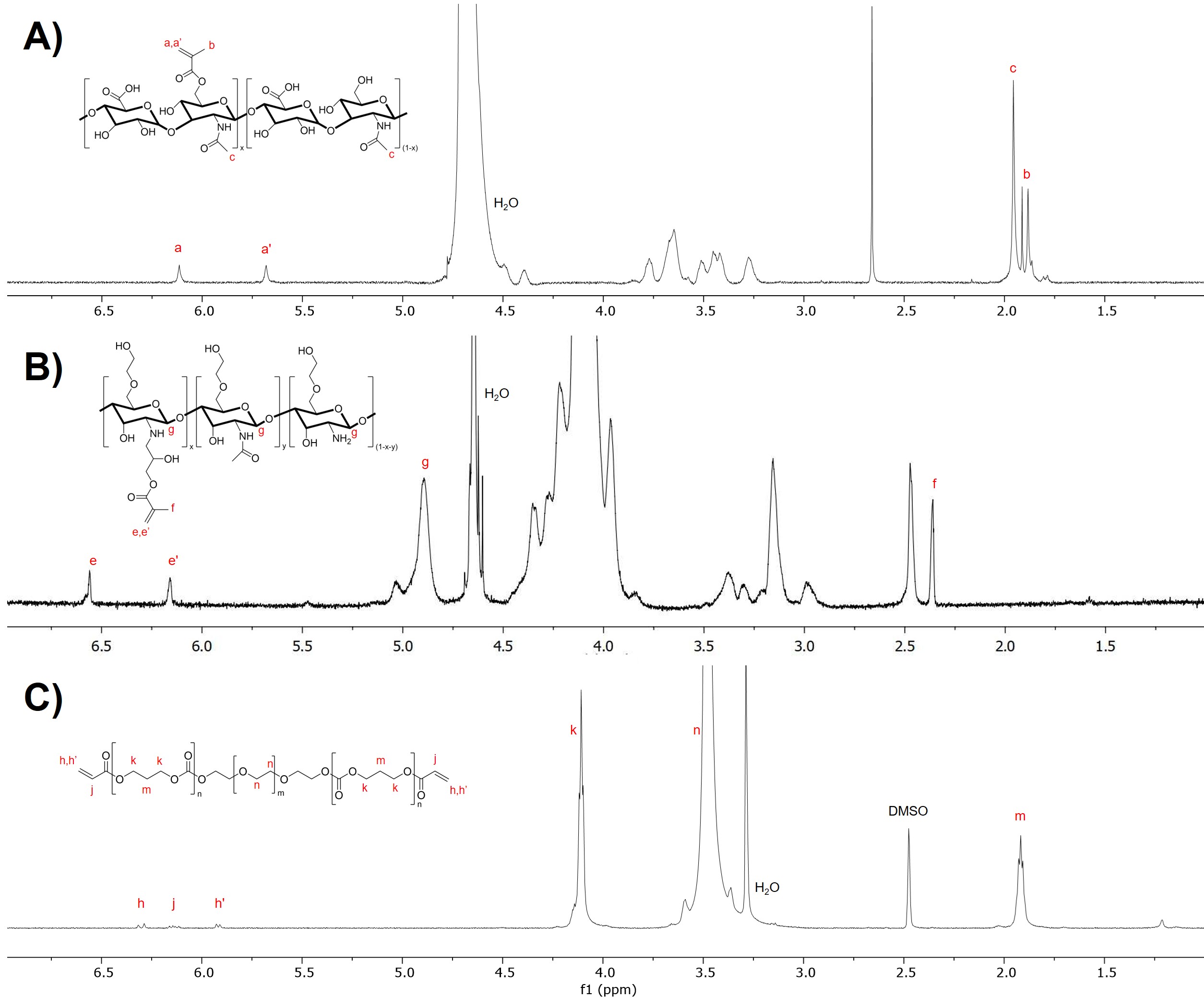

Supplementary Figure S1: ^1^H NMR was used to confirm the structure and degree of functionality of the polymers. (A) For MHA, the degree of methacrylation was calculated as ~25% from the ^1^H NMR spectrum. (B) For MGC, the degree of methacrylation was calculated as 5% from the ^1^H NMR spectrum. The fraction of residual acetyl groups was described by the supplier as 15%. (C) Molecular ratios of PEG_20_(PTMC_2_-A)_2_ were determined, and the degree of acrylation was 85%, based on the ^1^H NMR spectrum.
