## Supplementary Figure S3 for "Probing the effects of polysaccharide hydrogel composition on the viability and pro-angiogenic function of human adipose-derived stromal cells"

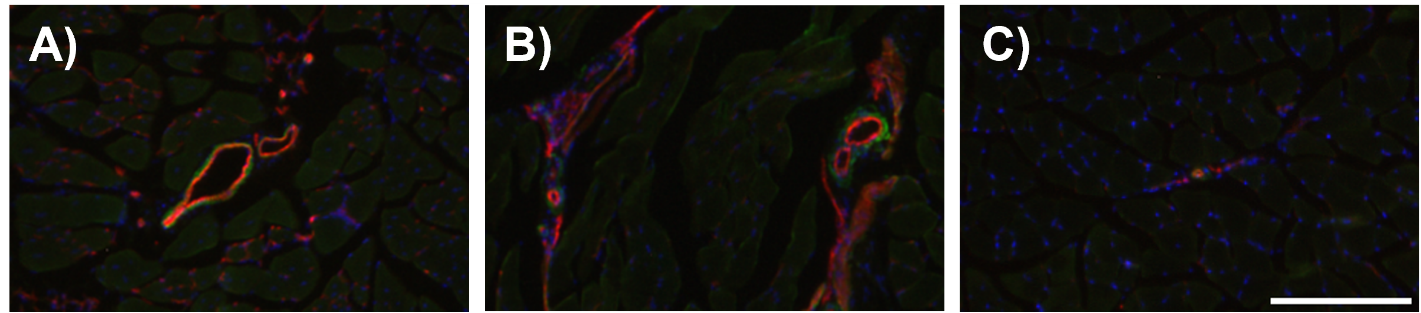

Supplementary Figure S3: Representative images of intramuscular (IM) vessels of different size ranges used in the scoring of IF samples within the gastrocnemius muscle harvested 28 days post-surgery, following FAL-CLI in athymic nude mice. (A) large IM vessels, (B) medium-sized IM vessels, (C) small IM vessels. CD31^+^ endothelial cells (red), α-SMA^+^ cells (green), DAPI^+^ cell nuclei (blue). Scale bar represents 200 μm. Scoring was assessed based on the size and frequency of vessels within the muscle, or within the region where FAL was performed.
