## Supplementary Figure S2 for "Probing the effects of polysaccharide hydrogel composition on the viability and pro-angiogenic function of human adipose-derived stromal cells"

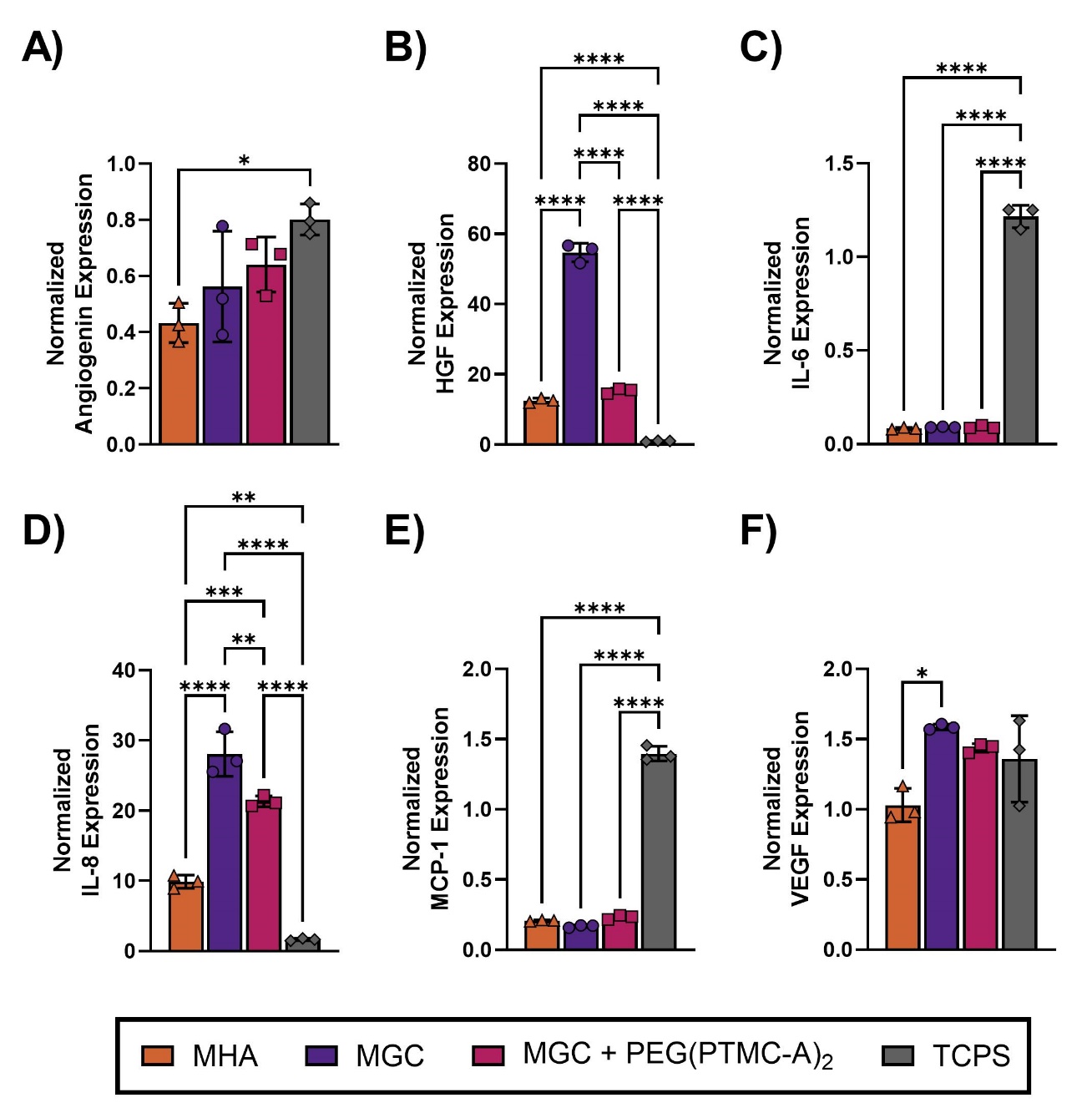

Supplementary Figure S2: Encapsulation within polysaccharide hydrogels of varying composition influences the hASC phenotype. Gene expression levels of angiogenin (A), HGF (B), IL-6 (C), IL-8 (D), MCP-1 (E), VEGF (F), isolated from hASCs encapsulated and cultured *in vitro* 7 days in the three hydrogel formulations or on tissue culture polystyrene (TCPS), normalized to IPO8, RPL13 and UBC housekeeping genes. All data presented as mean ± SD. Differences between hydrogel groups were determined using one-way ANOVA, with Tukey’s correction for multiple comparisons; Significant differences are indicated (* p<0.05, ** p<0.01, *** p<0.001, **** p <0.0001), with n=3 hydrogels for a single hASC donor.
